# Temporal dynamics of grooming reciprocity in crested macaques: Strongly bonded individuals reciprocate grooming more quickly

**DOI:** 10.64898/2026.09.21.753130

**Authors:** Luke Collins, Adam Gaston, Juliane Kaminski, Jerome Micheletta, Marine Joly

## Abstract

Primates maintain cooperative relationships largely through grooming reciprocity, yet there is no agreement on the timescales over which grooming is returned. Part of this disagreement stems from statistical issues: inconsistent definitions of a reciprocation window, combined with the exclusion of unreciprocated grooming, leave the debate unresolved. We still do not know if primates exchange grooming within the short windows predicted by biological market theory or over much longer timescales. Whilst survival analysis can identify reciprocation windows objectively by retaining unreciprocated grooming as censored data, previous studies have applied it inconsistently. This inconsistency leaves the true time frames over which primates reciprocate grooming somewhat blurred. Here, we use a statistical approach that addresses these problems to examine grooming reciprocity in crested macaques (*Macaca nigra*), a socially tolerant macaque species. We applied survival analysis to identify each group’s short-term reciprocation window without imposing arbitrary cut-offs, and Bayesian survival models to test whether social bond strength and eigenvector centrality predicted reciprocation latency. Short-term reciprocation occurred rarely (29% of reciprocal events in one group and 5% in the other); most reciprocation occurred over longer timescales than previous research commonly assumes. Stronger bonds predicted faster reciprocation; this finding suggests that crested macaques express social investment in valued partners through how promptly they return grooming, meaning that the timing of exchange carries information about relationship value. Eigenvector centrality showed no credible effect, though our small sample size and dense groups prevent us from interpreting this outcome more robustly. Substantial variation between dyads remained unexplained, pointing to partner-specific properties we have yet to identify. These findings caution against assuming that reciprocity operates over short windows. They show that social predictors, such as bond strength, remain essential to understanding when primates return grooming. This work provides a baseline for testing whether grooming is exchanged for other social commodities, and for comparing reciprocation timescales across macaque species that differ in social tolerance.

## Introduction

Grooming is one of the most ubiquitous cooperative behaviours in group-living primates. It functions as a primary currency of social exchange and a key mechanism through which relationships are formed and maintained (Schino & Aureli, 2010; Silk, 2007a). Biological market theory (BMT) provides a framework for understanding why grooming is exchanged. Originally an economic model, BMT applies the principles of supply and demand to animal behaviour, framing social transactions of commodities between individuals (Bowles & Hammerstein, 2003; Noë & Hammerstein, 1995; 1994). A core prediction of BMT is that individuals should preferentially direct grooming toward those partners who reciprocate most. Partner choice is central to social interactions in BMT where at the proximate level, two individuals can trade social commodities and receive mutual benefits (Schino & Aureli, 2017; 2010). However, competition may subsist with potentially preferable partners elsewhere (Noë & Voelkl, 2013). Importantly, this does not require a cognitively demanding mechanism like calculated reciprocity, rarely seen outside of humans and requiring partner evaluation of future benefits (Schino & Aureli, 2017; Brosnan & de Waal, 2002). BMT suggests that cooperation can be sustained through attitudinal reciprocity, or emotional bookkeeping, between partners (Schino & Aureli, 2017; Brosnan & de Waal, 2002). Schino and Aureli (2009) argue that reciprocity is maintained through ‘emotional bookkeeping’, a mechanism driven by recent partner-specific interactions. Individuals track the emotional quality of an interaction rather than tracking and comparing past exchanges, which makes it less cognitively demanding than calculated reciprocity (Schino & Aureli, 2017; Brosnan et al., 2010). Positive interactions boost a partner’s perceived value and improve the likelihood of future cooperative behaviour through accumulated emotional states (Schino & Aureli, 2009; 2017). Consequently, reciprocity can occur over a longer time frame because the emotional record can persist and update gradually (Schino & Aureli, 2017; Brosnan et al., 2010).

Evidence for preferential long-term partner choice consistent with BMT predictions exists across a range of taxa. For example, controlled experimentation in vampire bats (*Desmodus rotundus*) showed bats that fed more non-kin in previous years received food donations from a larger network of non-kin bats in return (Carter & Wilkinson, 2015). Grooming reciprocation has similarly been documented across a wide range of primate species, yet questions remain over the time frame over which exchanges occur as they vary substantially. Clarifying this temporal dimension is important on two fronts. Firstly, it evaluates whether observed exchange patterns align with BMT predictions. Secondly, distinguishing between short- and long-term reciprocation reveals how far primates can extend the cognitive tracking involved. Maintaining reciprocity over a longer delay demands more of that capacity than an immediate exchange does (Sánchez-Amaro & Amici, 2015). Temporal discounting compounds this demand: Stevens and Hauser (2004) define temporal discounting as the devaluing of future rewards, which often produces a preference for smaller, immediate rewards over larger, delayed ones. Any study attempting to draw conclusions about grooming reciprocation therefore depends on how it measures the time frame of reciprocation (Sánchez-Amaro & Amici, 2015).

Previous analyses assessing primate grooming reciprocation time frames have suffered from key methodological limitations, including i) the varying cut-off points by which authors have defined a single grooming bout as ceased, and ii) the abundance of unilateral grooming interactions spread across grooming bouts and their removal from analyses (Dunayer & Berman, 2016; Sánchez-Amaro & Amici, 2015). Grooming bout cut-off points can range from 5–60 seconds depending on the study (Dell’Anna et al., 2024; Xia et al., 2013; 2012; Frank & Silk, 2009; Chancellor & Isbell, 2009; Gumert & Ho, 2008; Schino et al., 2003). This is problematic when inferring time windows of reciprocation and making species evaluations: comparisons between survival analysis and original cut-off points used in previous studies revealed that shorter definitions of reciprocal time-matching significantly underestimated the actual rates of reciprocity (Dunayer et al., 2019; Sánchez-Amaro & Amici, 2015). Whilst cut-off points for single grooming bouts may be sufficient to capture some reciprocation, reciprocation could similarly be missed. This means that conclusions about whether and how quickly grooming is reciprocated are partly an artefact of methodological approach rather than a reflection of the underlying behaviour (Dunayer et al., 2019). The varying focal observation lengths additionally compound statistical issues further. For instance, reciprocation captured within a 30-minute focal may be missed entirely in a shorter observation window (Manson et al., 2004; Xia et al., 2012). Cross-study comparisons of time frames could be of limited validity as a result. Furthermore, primates regularly exhibit unilateral grooming interactions, yet multiple studies have used only within-bout reciprocal exchanges for analyses (Sánchez-Amaro & Amici, 2015). For example, only 5-7% of total grooming bouts were immediately reciprocated in bonnet macaques (*Macaca radiata;* Manson et al., 2004). Reciprocal grooming exchanges were also observed over the intermediate term, not within single grooming bouts in capuchins (*Cebus apella*; Schino et al., 2009) and olive baboons (*Papio anubis*; Frank & Silk, 2009). These interactions are not failures of reciprocation: they are biologically relevant events that hold meaningful information about the full time course of exchange, and their exclusion introduces systematic bias toward apparently shorter reciprocation windows.

Survival analysis directly resolves both problems outlined above: it identifies the short-term reciprocation window objectively by comparing the rate of grooming reciprocation following a received grooming interaction against the group’s collective baseline grooming rate (Dunayer et al., 2019). It imposes no arbitrary cut-offs: grooming interactions that occurred within and across focal observations can be included, whilst simultaneously accommodating unreciprocated interactions as censored data rather than discarding them (Dunayer et al., 2019; Sánchez-Amaro & Amici, 2015; Tyrrell & Berman, 2012; Schino & Pellegrini, 2009; Schino et al., 2009). Despite these advantages, its use remains infrequent across the literature, but it has been used in some previous macaque and other primate studies (O’Hearn et al., 2022; Dunayer et al., 2019; Schino & Pellegrini, 2009; Schino et al., 2009). Whilst there is not one perfect methodological choice, these confounds highlight the need to use appropriate statistical analyses. To our knowledge, two previous studies have applied survival analysis to grooming reciprocation in crested macaques *(M. nigra)* which provide a comparative baseline for our study (Dunayer et al., 2019; Tyrrell & Berman, 2012).

Even within a single social group, individual variation in sociality shapes the number of partners individuals interact with, the frequency of those interactions, and the broader connectivity patterns of their network position (De Moor & Brent, 2025). Within-group variation in sociality should have profound effects on fitness (Silk, 2007b). Previous research on partner choice across primate species has demonstrated that stronger social bonds influence the time frames within which grooming is reciprocated (Surbeck & Hohmann, 2015; Cheney et al., 2010). Even dyads within the same group may exchange over different time frames (Sánchez-Amaro & Amici, 2015). However, the direction of this effect could theoretically be contested based on factors including species, cognitive ability, tolerance, or social system. On one hand, looser bonds could rely more upon contingency-based exchanges that provide predictability in the short-term, whilst dyads characterised by stronger bonds could show less reliance upon short-term contingencies, be more likely to tolerate unbalanced exchanges and reciprocate over longer time frames (De Moor & Brent, 2025; Surbeck & Hohmann, 2015; Jaeggi et al., 2013; Seyfarth & Cheney, 2012). Alternatively, strongly bonded dyads may reciprocate more quickly, actively investing in and maintaining valued partnerships through prompt return of grooming, consistent with evidence that time-matched grooming has been reciprocated within bouts in some species (Chancellor & Isbell, 2009; Barrett et al., 1999). Individual network centrality is a secondary social factor not commonly investigated for its effects on grooming reciprocation. Centrality measures can provide more social information beyond interactions between dyads and can reflect the degree to which individuals are embedded within the broader social network of the group (Schülke et al., 2022; Farine & Whitehead, 2015; Croft et al., 2008). Central individuals are important to group cohesion and display different social characteristics compared to less central individuals (Schülke et al., 2022; Brent, 2015). It seems reasonable to assume that individuals occupying more central positions could have different social strategies compared to those less central, and thus the cooperative behaviours they employ (Gokcekus et al., 2021), including how quickly they exchange grooming, may itself be shaped by their network position. Social network analysis offers a method to quantify consequences of individual social differences (Wascher et al., 2018). It currently seems to be unknown if network centrality has an effect on the latency of grooming reciprocation. Social factors and partner choice options could also structure reciprocation timing, and future work needs to assess them directly (Schino & Aureli, 2017, 2010; Sánchez-Amaro & Amici, 2015).

Macaque species vary substantially in their degree of social tolerance, arranged along a four-grade scale from despotic (grade-1: rhesus macaques, (*M. mulatta)*, or Japanese macaques, (*M. fuscata*)) to tolerant (grade-4: Tonkean macaques *(M. tonkeana)*, or crested macaque *(M. nigra)*, Thierry, 2007)). Crested macaques are an ideal system in which to investigate how social bond strength affects reciprocation latencies. Social interactions in tolerant species are less constrained by nepotism, and dominance hierarchy exerts a weaker influence on social living compared to more despotic macaque species (Duboscq et al., 2017; Thierry, 2007). Crested macaque groups specifically contain a larger network of average-strength bonds where stronger bonds are more equitable but less predictable than weaker bonds (Duboscq et al., 2017; 2013). This social fluidity gives opportunity to observe temporal-bond social dynamics that may not be observable in more hierarchical macaque societies more constrained by relatedness or rank (Call, 2004; Thierry, 2000). Centrality effects have also previously been found across four macaque species along the tolerance spectrum, with top-ranking individuals in despotic species more likely to occupy central positions, suggesting differences in social strategy across the gradient (Sueur et al., 2011), though whether centrality independently predicts reciprocation latency within a single tolerant species remains unknown. Macaque species sitting at the tolerant end of the scale are more likely to show a different set of cognitive abilities due to their social organisation (Call, 2004, p.33). They are also comparatively understudied relative to despotic macaque species making them an underrepresented social system for investigating these questions (Loyant et al., 2023; Thierry, 2022; 2000; Joly et al., 2017; Duboscq et al., 2013; Call, 2004, p.33).

Firstly, we tested if overall grooming given was contingent upon grooming received. We tested this across all dyads irrespective of relatedness or dominance hierarchy. Secondly, using survival analysis and Bayesian modelling, we aimed to i) assess the time frames for short versus long-term reciprocation without imposing arbitrary cut-off criteria, and ii) determine whether longer reciprocation durations correlated with stronger dyadic social bonds or centrality positions. Given that stronger bonds in crested macaques are more equitable but less predictable than weaker bonds (Duboscq et al., 2017), we predicted that stronger bonds would be associated with longer reciprocation latencies. The logic here being that the more fluid social style of crested macaques would be reflected in a tolerance of temporary imbalance supported by relationship security. We also predicted that higher centrality scores would be associated with faster reciprocation to reflect the different individual social strategies. Together, these analyses include social factors not yet previously examined in crested macaques to assess their effect on the temporal structure of their grooming reciprocation.

## Methods

### Ethical note

All research adhered to legal requirements and guidelines set by the Animal Welfare and Ethical Review Body at the University of Portsmouth (approval: AWERB-223A). Our research was additionally supported by the BIAZA research committee, UK.

### Subjects & housing conditions

We observed a total of 20 macaques belonging to two captive crested macaque groups housed at Zoo de CERZA (CERZ), France (N = 11; seven females, four males) and Emerald Park (EMER), Ireland (N = 9; five females, four males) between September 2023 and April 2024. All subjects had known maternal genealogies, and we obtained taxon reports for all macaques in both groups. Observer LC reliably distinguished between individuals by using facial and body features. Both zoos are accredited members of EAZA (European Association of Zoos and Aquaria) and follow their established animal welfare protocols; Emerald Park is additionally a member of BIAZA (British and Irish Association of Zoos and Aquariums). All macaques were familiar to human presence and had access to indoor and outdoor enclosures during observations. Indoor and outdoor enclosures at both institutions were equipped with climbing structures and contained water ad-libitum. No macaques were separated for our research and they had continual contact with other group members. CERZ had recently introduced a new breeding male into the group in April 2023. Both groups were fed in the morning and afternoon a mixture of fruits, vegetables, nuts and seeds, and monkey pellets scattered around their outdoor enclosures.

### Data collection

#### Social bond strengths

We collected a total of 150.5h of focal observations (CERZ: 76.5h, M = 7h per individual; EMER: 74h, M = 8h per individual). Social bond observations were 30-minute continuous focal samples recorded from the enclosure periphery at CERZ, and both outside and inside enclosures for the group at EMER. Our measure of social bond strength was calculated using matrices of grooming, affiliative, and proximity behaviours to create a single composite sociality index (CSI) score (Duboscq et al., 2017; Silk et al., 2013; Silk et al., 2006):

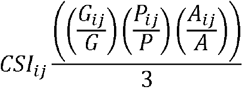

G*ij* is the grooming rate between individual *i* and *j* (duration of grooming given and received in min per hour of dyadic observation time) and G is the mean grooming duration across all dyads; P*ij* is the rate of proximity between individual *i* and *j* (frequency subjects were within one body length per hour of observation time) and P is the mean frequency of proximity for all dyads; A*ij* is the rate of affiliative physical contact between individual *i* and *j,* and A is the mean rate of affiliative contact for all dyads. Non-observed dyads for indices were assigned CSI = NA before averaging the final CSI score. Dyads with only one grooming interaction were removed from analyses. We decided to compute the grooming CSI score using duration over frequency for a more reliable measure. The mere occurrence of a grooming interaction (rather than duration) can be used to facilitate non-grooming social exchanges whereas duration represents invested grooming time to maintain the social bond (Sánchez-Amaro & Amici, 2015). Stronger social bonds were represented by a higher CSI score and lower CSI scores represented weaker social bonds.

We obtained data for all possible 55 dyads at CERZ (N = 11 individuals), and for all 36 possible dyads at EMER (N = 9 individuals). Focal subjects were randomly generated each day; upon the whole group being observed once, we repeated this cycle until data collection ended. We defined grooming as an individual combing the hair of another using the hands or mouth (Xia et al., 2021). Grooming behaviour was separated into two categories: *grooming duration* refers to the length of time an individual spent grooming a partner; and a *grooming episode* refers to a single continuous period in which one individual groomed another, ending when the groomer lifted its hand off and stopped grooming. Grooming start/end time was recorded along with identification of whether an individual was giving or receiving grooming. Every second minute we recorded proximity data of individuals within body contact, and 1 and 5 body lengths. Affiliative behaviours included touch, embraces, hugs, and lip smacks (Thierry, 2000).

### Social network analyses

We used the *igraph* R package (Csárdi & Nepusz, 2006) to create a matrix of grooming, affiliative and proximity interactions to measure degree, strength and eigenvector centrality. Degree and strength captured direct connections: the number of unique partners, and the summed weight of an individual’s interactions with them (Brent et al., 2011). Eigenvector centrality (EC) captured indirect position instead, weighting an individual’s connections by the centrality of their partners (Brent, 2015). We selected these three measures because each has separately been linked to neurocognitive outcomes in macaques. The number of an individual’s direct social partners predicts the volume of brain regions implicated in social decision-making in free-ranging rhesus macaques (Testard et al., 2022). Independently, EC in a grooming network is linked to inhibitory control task performance, though not to other cognitive domains, in wild Japanese macaques (Kaigaishi & Yamamoto, 2024). For each behavioural measure, dyadic frequencies were first normalised to a rate per minimum dyadic observation hour and then z-score standardised. The three z-scores were then averaged to produce a single score per dyad. To minimise overestimation of dyadic interaction opportunity by using the total sum of dyadic observation time, we used the minimum dyadic observation time to measure normalisation rates for z-scores. Using minimum was preferable as a more conservative proxy of dyadic interaction time and to help with the underestimation of strong dyads without inflating weak bonds (Davis et al., 2018; Farine, 2015).

### Dominance characteristics

Elo dominance ratings were calculated using the EloRating R package (Neumann et al., 2011) following the method originally proposed by Albers & de Vries (2001). Dominance positions were calculated by an initial starting rate (k = 100) and individual scores were updated by an amount dependent on individual A’s chance of winning against individual B. We used the dyadic maximum likelihood (DML) estimator by Milligan (2003) to quantify mother-child relatedness; we had 4 categories of relatedness across groups (grandparent, parent, siblings, uncle/aunt).

### Video analysis

All focal samples were videotaped and uploaded to BORIS (Behavioural Observation Research Interactive Software; Friard & Gamba, 2016) for behavioural coding. All behaviours along with macaque IDs were verbally confirmed live by LC. We recorded grooming duration from start to end, affiliative behaviours (lip smack, hug, embrace, touch), and proximity scan samples every 2 minutes (body contact, 1 and 5 metres).

### Statistical analyses

R was used for all analyses (R Core Team, 2024). A Tweedie generalised linear mixed model (GLMM) with log link using the glmmTMB package (Brooks et al., 2017) was used to establish if overall dyadic grooming given was dependent upon how much grooming was received. Grooming given and received for both groups were combined for analysis. The dyadic grooming given rate was our response variable and the rate of grooming received was our predictor with group as a fixed factor and dyad ID as a random factor. The grooming rates were calculated by using the total seconds of grooming given and received divided by the total dyadic observation time. Grooming rates were used over raw duration to control for differences in observation time and create a directly comparable rate of grooming. A Tweedie distribution was used as its function more easily accommodates near zero values and right skew distribution of data. For simplicity, our analyses purely focussed on grooming and no other type of inter-commodity exchange. Whilst grooming interactions have been shown to be traded for other social commodities, it is difficult to properly assess reciprocity where the social commodities are not the same (Brosnan & de Waal, 2002; Schino & Aureli, 2008b). Model assumptions and fit were checked using the DHARMa package (Hartig, 2022). We conducted QQ plots to verify uniformity of residuals; residual versus predicted values were inspected visually for homoscedasticity, and we ran non-parametric dispersion and zero inflation tests. We ran our full model against a null model that contained only the dyad ID random effect using a likelihood ratio test to confirm the predictor groom received explained some variance above chance.

#### Survival analysis

We identified short- and long-term reciprocation windows by survival analysis (package: muhaz, Hess & Gentleman, 2026) for an elapsed time window in which an original groomer was groomed back by their partner at a rate greater than baseline grooming rates (hazard rate; Dunayer et al., 2019). Time between the end of the grooming episode and the end of the focal session was entered as censored data. If grooming was not reciprocated within the focal observation, the elapsed time continued to accrue across subsequent focal observations of either dyad member until reciprocation occurred or the observation ended. Survival analysis caters for censored data suitably as focal observations can conclude without reciprocation occurring. A smooth hazard rate was plotted to estimate approximately how long it would take the groomee to return the previously received grooming. This was compared to the baseline rate of grooming. The baseline calculation is summed as the total sum of weighted rates / total number of grooming episodes (Schino et al., 2009), using the below calculation:

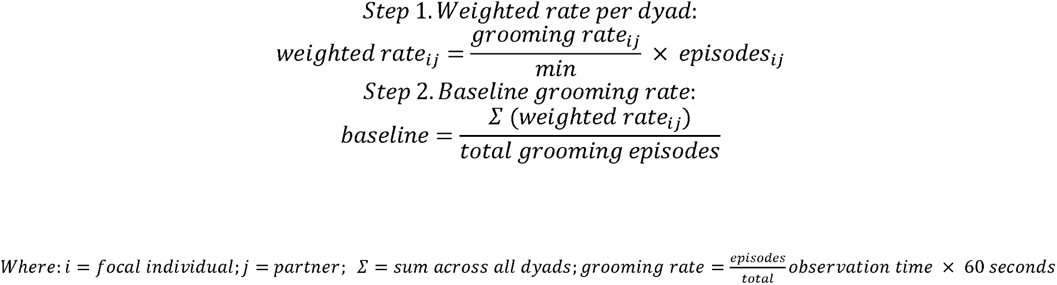

The baseline rate was plotted as a constant. Short versus long-term grooming reciprocation was determined at the point at which the hazard rate crossed the baseline rate, not the lower bound confidence interval as structured in previous studies (Dunayer et al., 2019). The reason for this was that EMER did not show a short-term window with the crossover point being placed at the lower bound of the confidence interval. Elapsed time windows outside of baseline rates were considered long-term grooming reciprocation, and time windows before the baseline rate were considered short-term reciprocation. We considered a grooming episode reciprocated when the groomee subsequently groomed the original groomer, i.e., a switch in the direction of grooming (Schino et al., 2009).

#### Bayesian survival analysis

To assess our main question of whether dyadic social bond strength affected reciprocation latency, we fitted two Bayesian survival log gamma models with a multi-membership term using the *brms* package (Bürkner, 2017). A model combining data from both groups was used for final analyses, and our fixed effects were z transformed. The first model included a response variable of combined reciprocation and censored durations into a single time column, with event coded as event = 1, and censored = 0, as required by the *cens* function in *brms* which expects both event times and censoring times in the same response variable. Social bond strength was the fixed effect. We did not include fixed effects of age, sex, relatedness or dominance rating as the model became unstable with additional covariates. This is to be expected in captive groups with high relatedness and a limited number of subjects. However, we did include dyad and group ID as random effects in an attempt to capture individual variance in our results. A multi-membership term (focal and partner ID) was included to account for non-independence of each individual within a dyad with equal weights assigned to focal and partner identity (brms default). A gamma likelihood distribution was used on account of right tail positive values. Priors were weakly informative on the intercept (log(195), 0.5) and informed by macaque grooming reciprocation literature that has used survival analyses to a mean of 195 seconds (O’Hearn et al., 2022; Dunayer et al., 2019; Majolo et al., 2012; Tyrrell & Berman, 2012). Gamma shape parameter was fit with (2,0.1) and a student-t (3,0,1) prior was placed on the random effects with adjusted initial values to ensure stable sampling. The second model consisted of the additional fixed effect of eigenvector centrality. Four chains were run for 4000 iterations each (2000 warm up), with adapt delta = 0.999 and max treedepth = 15 to minimise divergences on both models. Convergence was assessed via Rhat values, and bulk and tail effective sample sizes (>1000). Our analyses included reciprocal events both within and across focal observations, as restricting events to within a focal observation would have excluded longer reciprocation latencies (see Sánchez-Amaro & Amici, 2015 for full review of statistical issues with current grooming reciprocation analyses). We compared likelihood distribution fit using LOO (Leave-One-Out) cross validation and posterior prediction checks of density overlays (observed versus predicted posterior draws), mean and median predictions as per BARG (Bayesian Analysis Reporting Guidelines) (Kruschke, 2021).

## Results

For both groups, we observed a total of 1,892 grooming episodes (CERZ: N = 863, mean (M) = 1.17, standard deviation (SD) = 1.25 episodes/hour; EMER: N = 1029, M =1.77, SD = 1.64 episodes/hour), 2,192 affiliative interactions (CERZ: N = 1322, M = 1.8, SD = 2.05 episodes/hour; EMER: N =870, M = 1.57, SD = 1.25 episodes/hour), and 3,610 proximity interactions (CERZ: N = 1560, M = 2, SD = 1.27 episodes/hour; EMER: N = 2050, M = 3.46, SD 1.31 episodes/hour). CERZ social bond strengths ranged from 0.05 to 4.15 (M = 0.89, SD = 0.8) and EMER CSI strengths ranged from 0.1 – 2.82 (M = 0.94, SD = 0.58). Full dyadic CSI scores can be found in Supplementary Table S1.

### Is grooming given contingent upon grooming received?

To answer our first question of whether grooming given was dependent upon the grooming received, we analysed a total of 40,013 seconds (11.1 hours) of grooming given and 36,800 seconds (10.2 hours) of grooming received across both groups in 86 dyads (CERZ: 51 dyads, and EMER: 35 dyads). Dyadic grooming given was significantly dependent upon the duration of grooming received (β = 0.017, z = 5.93, p < 0.001 – see Supplementary Table S2 for full summary). CERZ showed a lower mean grooming given and received rate (CERZ: give = 19.21 ± 23.94, receive = 24.85 ± 32.43 seconds/hour) compared to EMER (EMER: give = 44.85 ± 46.22, receive = 31.90 ± 42.42 seconds/hour). EMER dyads showed a higher rate of giving grooming independent of how much grooming they received (β = 0.743, SE = 0.224, z = 3.33, p < 0.001). The full model provided a significantly better fit compared to the null model (χ² = 43.29, df = 2, p < 0.001; AIC full model = 714.3, AIC null model = 753.6). Additionally, the full Tweedie model with grooming rate as the response variable (AIC = 714) was a significantly better fit than the model using raw durations (AIC = 1154). DHARMa QQ plots showed no significant deviation from the expected distribution (KS test: p = 0.999), the dispersion test confirmed no overdispersion (dispersion ratio = 1.67, p = 0.27), and zero values were appropriately handled by the model (p = 1.00).

### Survival analysis

A total of 86 reciprocal grooming dyads (CERZ = 51 dyads, EMER = 35 dyads) were analysed in separate survival analyses. Survival analysis showed that the rate of reciprocated grooming was higher than the baseline group rates for 3 minutes for CERZ, and 34 seconds for EMER (see figure 1). Grooming reciprocation exchanged before these durations were considered short-term. Approximately 29% of CERZ (N = 47) reciprocal events were considered short-term reciprocation, and 5% (N = 8) for EMER. Mean reciprocation latency for CERZ was 26 minutes (SD = 31m), and 33 minutes for EMER (SD = 35m). More reciprocation occurred in a longer-term window for EMER than CERZ as the hazard rate for EMER drops below baseline at a quicker rate. For analysis compatibility, an upper limit of 180 minutes was applied to both reciprocation and censored duration outliers as this captured the scope of the data. One reciprocal event was removed from CERZ (N = 158 reciprocal events, N = 159 censored events), and 22 censored events were removed from EMER (N = 169 reciprocal events, N = 111 censored events). Sex differences were found in reciprocation latencies in CERZ with females reciprocating faster than males (log-rank test, p < 0.0001; see Supplementary Figure S1). The probability females would reciprocate past 100 minutes reached zero likelihood whilst male reciprocation outside of this time window did not occur. No significant sex difference in reciprocation latency was found in EMER (log-rank test, p = 0.32).

**Figure 1.**
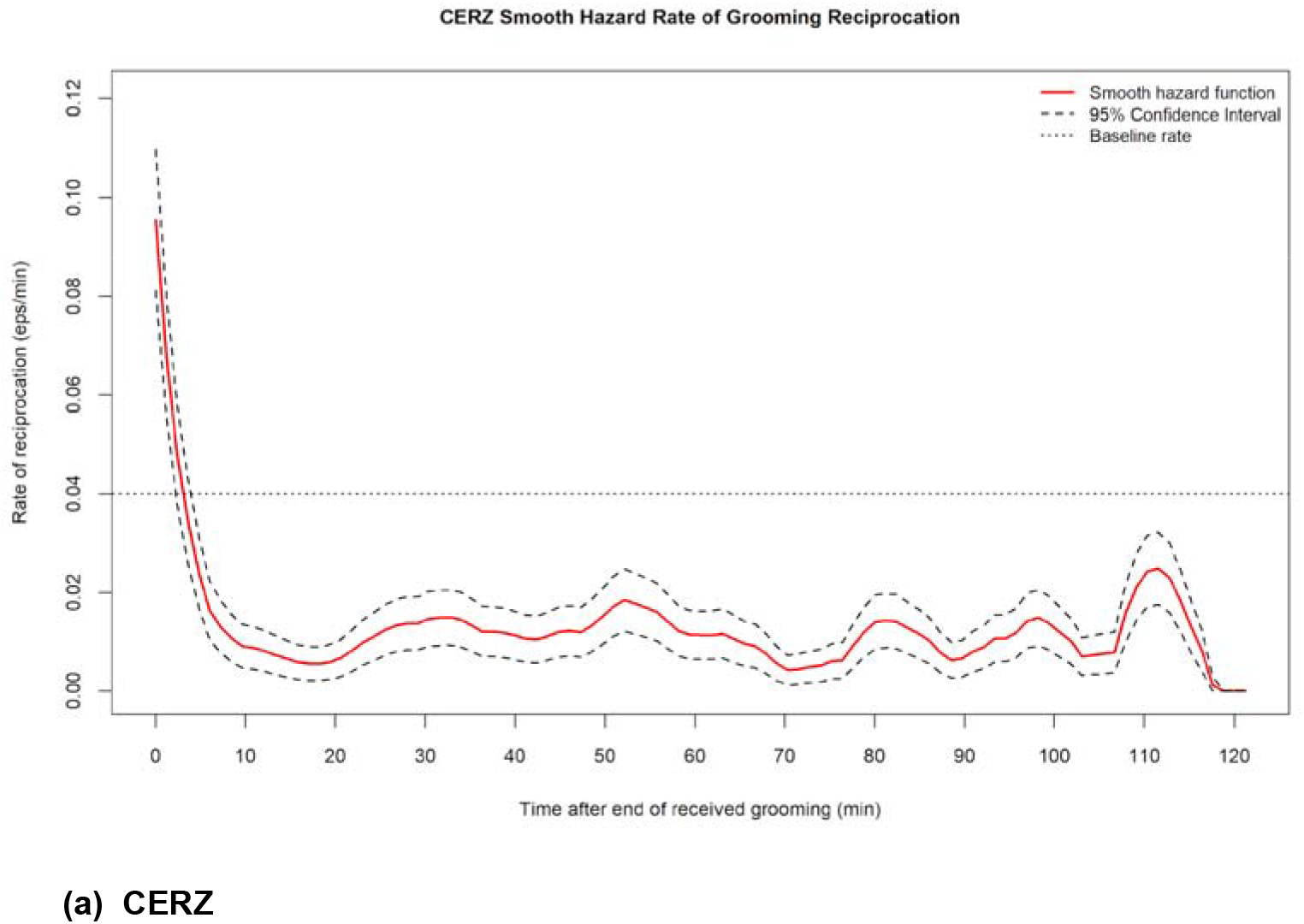
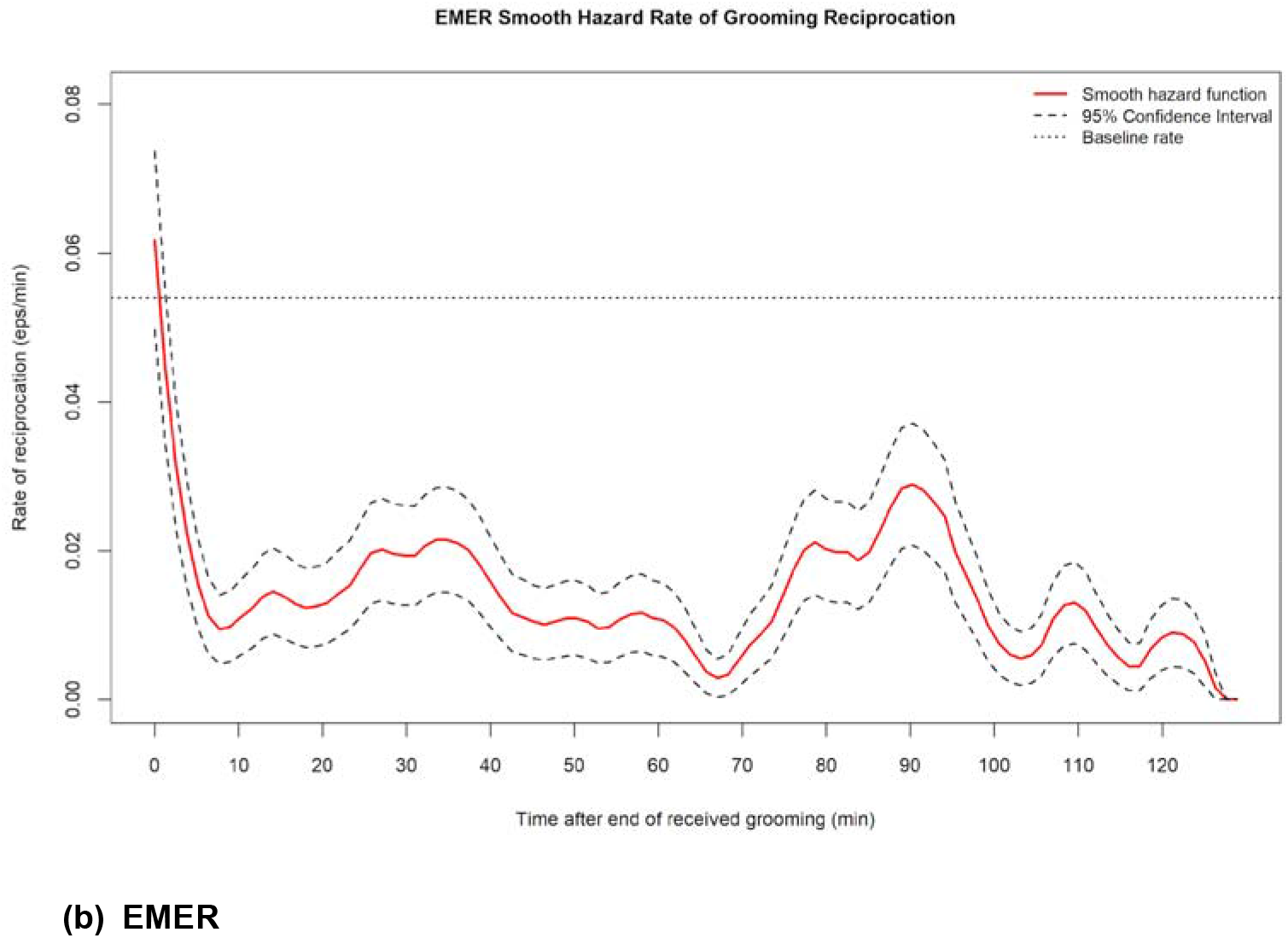
Smooth hazard rate of grooming reciprocation in CERZ and EMER. (a) CERZ (N = 158 reciprocal events) and (b) EMER (N = 169 reciprocal events). Solid red lines show the smooth hazard function and dashed lines show the 95% confidence intervals. Dotted horizontal lines mark each group’s baseline grooming rate (CERZ = 0.04, EMER = 0.05 episodes per minute).

### Bayesian survival analyses

Whilst we did not find evidence for stronger social bonds tolerating longer durations for reciprocation, we did find that more strongly bonded dyads reciprocated more quickly than weakly bonded dyads (β = -0.38, 95% credible interval (CI) [-0.67, -0.11] – see figure 2). The random effects of dyad ID, individual ID, and group explained some of the variance for latency to reciprocate. Dyad identity explained considerable variation in reciprocation latency (SD = 0.96 [0.67, 1.31]) with group membership contributing to the variation also but with wider credible intervals (SD = 2.58 [1.15, 5.57]). These results show that reciprocation latency was heavily influenced by dyadic and individual level characteristics as well as group-level differences (see Supplementary Table S3 for full summary). Some variation was captured by the multi-membership term but to a lesser extent, as the wide credible intervals suggest uncertainty (SD = 0.56 [0.03, 1.30]). The gamma likelihood distribution was tested against a hurdle lognormal full and null model. Posterior checks confirmed the gamma density overlay of observed versus predicted posterior draws more closely tracked the initial peak of reciprocation latencies and then the right tail decay compared to the hurdle lognormal model (see Supplementary Figure S2). Comparison of LOO cross validation showed greater gamma model accuracy compared to the hurdle lognormal model (difference in expected log predictive density (ELPD) = 34.9, SE = 8.5), with the hurdle lognormal full model only performing marginally better compared to a null model (ELPD difference = -0.9, SE = 8.5). All parameters achieved successful convergence across all four chains (Rhat = 1.00). Bulk and tail effective sample sizes exceeded the recommended threshold of 1000 for all parameters (Bulk ESS range: 1189 — 14669; Tail ESS range: 3073 — 6016). No divergent transitions were recorded.

**Figure 2.**
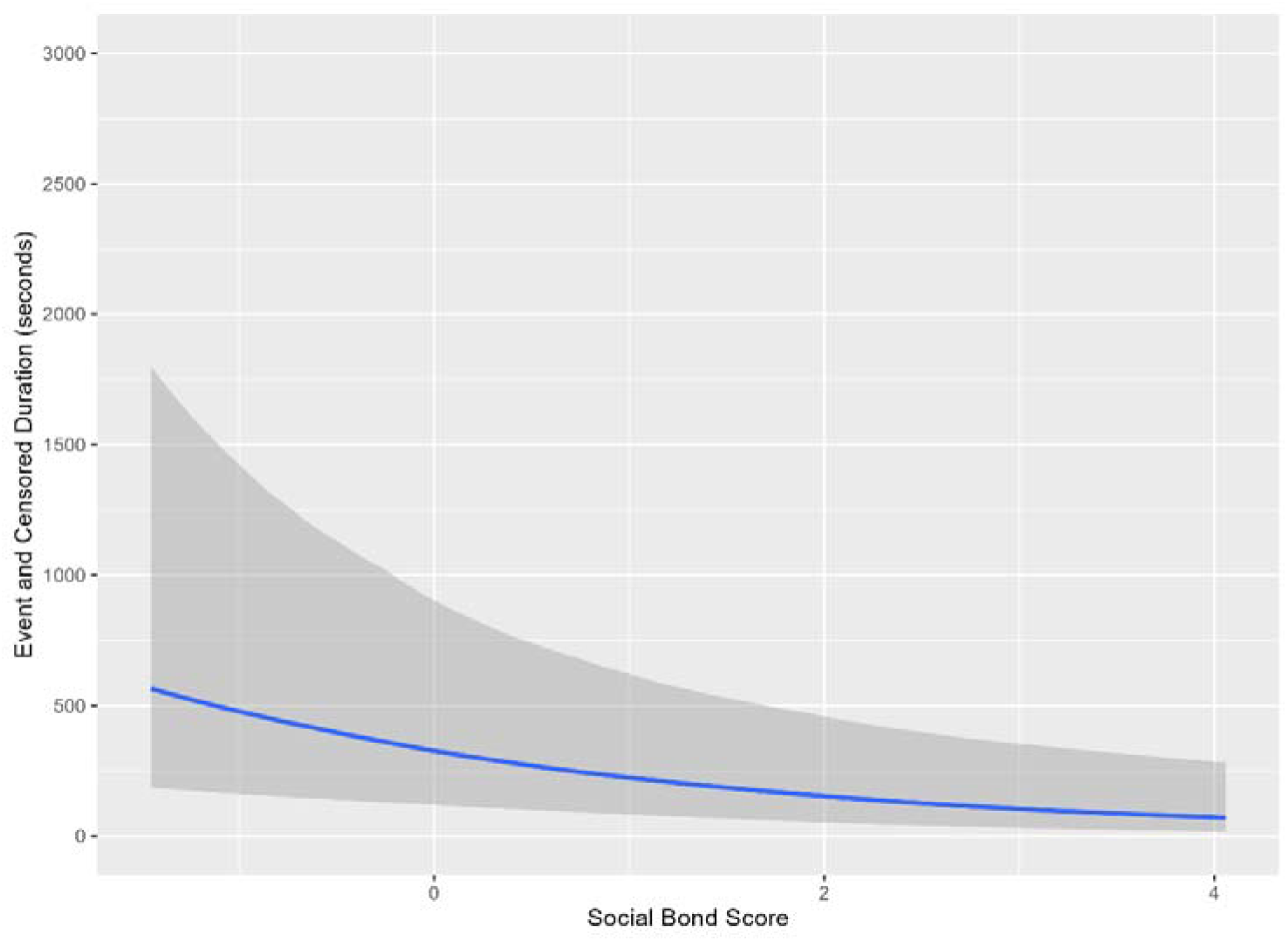
Predicted reciprocation latency across social bond strength. Predictions come from the Bayesian gamma survival model. The blue line shows the posterior median and the shaded area the 95% credible interval (N = 597 reciprocal and censored events from 86 dyads across CERZ and EMER). As social bond strength increases, the time it takes to reciprocate decreases.

We further investigated if individual level eigenvector centrality could explain some of the observed differences (see Supplementary Table S4 for centrality scores). We additionally included eigenvector as a fixed effect alongside social bond strength, but we did not observe a credible effect on reciprocation latency (β = 0.05, 95% CI [-0.25, 0.36]). The social bond strength credible effects remained aligned with our first model to confirm that observed reciprocation latencies were driven by dyadic bond strength compared to individual characteristics (β = -0.40, 95% CI [-0.69, -0.12]; see Supplementary Table S5 for full summary results). LOO cross validation confirmed that adding eigenvector provided no improvement compared to the sole social bond strength model (ELPD difference = -0.1, SE = 0.3; see Supplementary Table S6). However, the random effects were similar to the first model in that dyad identity (SD = 0.98 [0.67, 1.33]) and group identity (SD = 2.58 [1.12, 5.66]) contributed much to the variance with group identity exhibiting particularly wide credible intervals. The multi-membership term (SD = 0.58 [0.03 – 1.35]) again captured some variance but to a lesser extent as the lower CI was closer to zero compared to dyadic and group identity. Posterior predictive checks confirmed that the overlay of observed versus predicted replications strongly aligned with the sole social bond strength model and captured the gamma distribution of the model well (see Supplementary Figure S3). The predicted replication tracked the initial peak but still slightly underestimated the frequency of quicker reciprocation and overestimated right tail reciprocations. All parameters achieved successful convergence across all four chains (Rhat = 1.00). Bulk and tail effective sample sizes were confirmed for all parameters (Bulk ESS range: 1001 – 12674; Tail ESS range: 2572 – 5978). No divergent transitions were recorded.

## Discussion

Here, our main aim was to help contribute to a standardised methodological approach for analysing the time frames over which grooming reciprocation occurs in primate groups. Firstly, we wanted to establish an empirical baseline of what counts as short versus long-term grooming latencies in an understudied socially tolerant macaque, and secondly, to assess social factors affecting the timing of reciprocal grooming. We initially tested if grooming given was dependent upon grooming received across 86 dyads and found that dyadic grooming given was significantly dependent upon the duration of grooming received. Dyads that received more grooming gave greater grooming in return. Then, using survival analyses, we analysed reciprocation intervals that captured the full temporal grooming distribution without imposing arbitrary cut-off points and included non-reciprocated grooming episodes as censored data. We showed that reciprocation prior to 3 minutes for CERZ and 34 seconds for EMER was considered short-term grooming. Social bond strength was then used as a predictor to assess if this would affect grooming reciprocation latencies. Bayesian survival analyses showed that dyads characterised by stronger social bonds reciprocated grooming more quickly. Female individuals in CERZ were also observed to reciprocate grooming more quickly than males. Similarly, we wanted to assess if eigenvector centrality, a measure of how well connected an individual’s partners are, would help predict reciprocation latencies; however, we did not find any evidence for this.

### Grooming given was contingent upon grooming received

Our initial finding that dyadic grooming is preferentially given to those that have groomed them the most aligns with previous works across different primate species, including marmosets, capuchins, mandrills, baboons, and chimpanzees (Campennì et al., 2015; Surbeck & Hohmann, 2015; Frank & Silk, 2009; Schino & Pellegrini, 2009; Schino & Aureli, 2008a). Crested macaques, positioned at the tolerant end of the macaque social styles, show this same preferential grooming pattern as other macaque species (Roubová et al., 2015; Majolo et al., 2012; Xia et al., 2012; Gumert & Ho, 2008; Schino et al., 2007; Schino et al., 2003; Manson et al., 2004). This initial result provides a baseline for confirming that both groups have a dyadic contingency when directing their grooming. Partner choice is a core tenet of BMT (Noë & Hammerstein, 1994). A preferential dyadic contingency is consistent with several proposed mechanisms, including emotional bookkeeping of different partners and their previous interactions (Schino & Aureli, 2009). EMER was found to have a higher baseline rate of grooming given independent of how much grooming they received. One contributing factor for this may be the higher relatedness in this group. Kin-bias affiliation is well known in primate groups (Seyfarth & Cheney, 2012). In macaques, more closely related individuals have a higher probability of elevated affiliation rates and are more likely to tolerate asymmetries in grooming exchanges (Thierry, 2007; Chapais & Bélisle, 2004; Chapais, 2001). Similarly, this could also be a result of genuine group-level differences in social structure, but a larger number of groups would be necessary to assess this.

### Survival analysis

Having initially observed that dyadic reciprocity exists within the groups, we used survival analysis to determine the structure of reciprocation without imposing arbitrary temporal cut-offs, i.e., short versus long-term reciprocation. For CERZ, short-term reciprocity took place before 3 minutes and 34 seconds for EMER. Most reciprocal events occurred over longer-term time windows, with 71% of CERZ, and 95% of EMER reciprocal events considered long-term. The mean reciprocation time was 26 minutes for CERZ and 33 minutes for EMER with reciprocation occurring within and across focal observations. It is important to start by establishing the time frame over which reciprocation occurs and addressing the strengths and weaknesses of previously used statistical approaches before inferring results or assessing any BMT predictions (Sánchez-Amaro & Amici, 2015). The first is a statistical concern over defining what short-term reciprocation is. It appears that prominent issues with previous primate grooming analyses have been the variation of time frames being used to define short-term grooming and only including grooming bouts in analyses that were reciprocated (Dunayer et al., 2019; Sánchez-Amaro & Amici, 2015). Criteria to differentiate between short-and long-term reciprocation have often not been described; instead, definitions have been based on methodological consistencies with previous studies, identified via discontinuities in the frequency distribution of pause lengths, or chosen based on pause lengths that exceed the vast majority of pauses (Sánchez-Amaro & Amici, 2015). Existing literature tells us that the end of grooming bouts could be when grooming is suspended for 10 seconds, (Henzi & Barrett, 2002; Barrett et al., 1999), 20 seconds, (Fruteau et al., 2009), 30 seconds (Fruteau et al., 2011; Gumert, 2007), and longer durations still (Chancellor & Isbell, 2009; Port et al., 2009; Payne et al., 2003). Such variations in definitions would affect the grooming information being analysed and consequently the results (Sánchez-Amaro & Amici, 2015). The effect of arbitrary cut off points on analyses has previously been demonstrated, with shorter cut off points underestimating the degree of grooming time matching and influence of rank distance between dyads of rhesus macaques (Dunayer et al., 2019). To our knowledge, two previous studies have used survival analyses to assess grooming reciprocation in crested macaques (Dunayer et al., 2019; Tyrrell & Berman, 2012). Whilst both our analyses implemented a survival framework, Dunayer et al. (2019) applied a different statistical approach to the survival curve that asked a distinct question regarding what effect varying arbitrary cut-off points used by different authors had on short-term time matching. When they applied a slightly modified version of the analysis that included all censored events, they found short-term reciprocation occurred within 5.5 minutes (C. Berman, personal communication, April 28^th^, 2026; Tyrrell & Berman, 2012), which more closely aligns with our CERZ group. From the studies available, it seems more despotic species appear to have shorter time windows for short-term reciprocity. Cox survival analyses conducted on two macaque species with intermediate degrees of social tolerance showed the timing of reciprocity to be between five seconds and one minute (barbary macaques (grade-3): Molesti & Majolo, 2017; long-tailed macaques (grade-2): Majolo et al., 2012). Although some evidence exists for bonnet macaques (grade-3) males less likely to groom reciprocally within a day compared to a two-month period, but the methodological approaches are difficult to compare (Adiseshan et al., 2011). Our EMER duration of 34 seconds is more closely aligned with these long-tailed and Barbary macaque durations that resemble shorter and more direct reciprocation. However, the EMER hazard curve shows a more uncertain and rapidly dropping rate below EMER baseline rates of grooming which suggests dyads showed no real meaningful short-term reciprocation window at shorter durations. Moreover, we had to apply an upper limit of 180 minutes to censored events as including outliers above this threshold made it difficult to identify the point in time where baseline rates crossed over the hazard rate. If we retained those outliers, the hazard curve did not cross the baseline at any identifiable point. This could reflect genuine group dynamics because individuals share a closer genetic relationship in EMER compared to CERZ. Kin derive indirect fitness benefit from helping one another (Hamilton, 1964): relatives may not need to promptly return grooming. Kinship bonds also remain more stable and predictable across years than bonds between non-kin (Cheng et al., 2026).

#### Methodological influences on the observed timescales

Our own methodological approach could also be a contributing factor as each focal observation lasted 30 minutes, and as focal observation identity was randomly generated, a groomed individual in focal observation A was not necessarily the focal ID in focal observation B, and reciprocation could have easily been missed until it was captured later. It is also difficult to compare these studies within macaque literature and the broader primate literature. Methodological differences arise, further complicated by socioecological, housing and spatial proximity differences, or steepness of dominance hierarchy, which are all contributing factors that could affect reciprocity. More socially tolerant macaque species are less constrained by rank and nepotism (Duboscq et al., 2013; Thierry, 2007). Our short-term window for CERZ was much longer than reported for long-tailed or Barbary macaques (Molesti & Majolo, 2017; Majolo et al., 2012). Moreover, our data for EMER were uncertain for reciprocation rates at shorter time windows with a very large proportion of reciprocal exchanges even occurring up to 180 minutes for both groups.

It would be interesting in the future to assess temporal patterns of grooming reciprocation that reflect the different social styles of macaque species. It may be the case that more socially tolerant macaques maintain short-term exchange over longer time windows because bond maintenance could rely less upon immediate contingency. More socially tolerant macaques have already shown different cognitive abilities represented through their different social tolerance styles (Duboscq et al., 2017), impulsive reactions (Loyant et al., 2023), and social cognition abilities (Bhattacharjee et al., 2026; Joly et al., 2017). The generalisability of future studies should have a statistical focus surrounding the use of survival analysis to objectively capture what time frames primate species reciprocate over (Dunayer et al., 2019; Sánchez-Amaro & Amici, 2015). Establishing the time frame over which grooming-to-grooming reciprocations occur provides an empirical baseline for evaluating the cognitive mechanisms that sustain them. Immediate contingency based mechanisms such as tit-for-tat reciprocity are sufficient for shorter-term exchanges, whereas reciprocation across extended periods would require more reliance upon mechanisms such as emotional bookkeeping (Schino & Aureli, 2009). Some of the extended mean reciprocation latencies we observed here represent this. Reciprocity then becomes more complicated when incorporating additional factors such as different social commodities, partner value, or seasonal fluctuations (Schino & Aureli, 2017; Schino & Pellegrini, 2009; Schino et al., 2009). A robust empirical baseline of grooming reciprocation is a prerequisite for assessing the cognitive demands placed on individuals and disentangling the exchange patterns and the time frames in which they occur.

A second statistical concern is including only reciprocal grooming bouts that occurred in shorter time frames and excluding unilateral grooming episodes. One-sided grooming interactions regularly occur in primate grooming. Excluding them is problematic because unilateral grooming exchanges are not failures of reciprocation. They are biologically relevant aspects of grooming strategies that hold important information about primate exchanges, which naturally elongate reciprocal durations of dyadic grooming exchanges (O’Hearn et al., 2022; Majolo et al., 2012; Schino et al., 2009; Schino & Pellegrini, 2009). Primate literature has generally considered reciprocal primate grooming to occur over shorter time periods because of the cognitive abilities required, such as the cognitive load of exchanging varying social commodities with multiple partners, and the natural inclination for temporal discounting with primates more likely to return a service immediately than later (Barrett & Henzi, 2006). Despite this assumption, there is plentiful evidence of cooperative long-lasting relationships in primates (Seyfarth & Cheney, 2012; Silk et al., 2010; Mitani, 2009; Silk, 2002). Our results are consistent with previous primate work that has shown reciprocation can occur over intermediate or longer time scales and challenges BMT predictions that grooming should be exchanged within single bouts (Barrett et al., 1999). For example, we observed a large proportion of unreciprocated grooming episodes entered as censored data (CERZ: N = 159, EMER: N = 111). Similar studies have shown that immediately reciprocated bouts in bonnet macaques accounted for only 5-7% of total grooming observed (Manson et al., 2004), and that grooming bouts became more equitably balanced between partners across multiple bouts rather than single bouts (Frank & Silk, 2009). Focal sampling methods will always pose inherent observational constraints, which is why it is relevant to include unilateral grooming episodes and treat them as censored rather than absent entirely. The results obtained in both groups in this study reflect the broader temporal window over which grooming occurs, based on the high volume of censored observations and extended mean reciprocation latencies. This further raises the question of dyadic level factors predicting these temporal windows and the influence of social bond strength accelerating or delaying reciprocity.

#### Social bond effects on reciprocation

Our main finding was that stronger social bonds predicted shorter reciprocation latencies. This suggests that in our groups of crested macaques, individuals apply a temporal contingency to their reciprocation and return grooming faster to those that they have a stronger bond with. This pattern is consistent with partner choice models of reciprocity in BMT where individuals preferentially direct cooperative effort towards partners they deem to be high value (Schino & Aureli, 2017; Campennì & Schino, 2014). These results extend the perspective of partner choice by showing that bond strength predicts how quickly grooming reciprocation occurs, indicating that the timing of reciprocity itself is temporally structured in crested macaques. This aspect adds a temporal dimension to maintaining social bonds where even dyads within the same group and species exchange over different time frames, often missing from broader conversations of reciprocity (Sánchez-Amaro & Amici, 2015). This finding contrasts with our initial hypothesis that stronger bonds would show longer reciprocation latencies. Our idea stemmed from established ideas of partner choice flexibility and emotional mediation theory. If individuals prefer to choose their partners based on competing bids of value, one possibility is that the bond is maintained even if the reciprocity is delayed because of attached longer-term fitness benefits (Jaeggi et al., 2013; Seyfarth & Cheney, 2012). For example, such a pattern has been reported in bonobos where individuals who were more familiar with each other and associated more had longer reciprocation latencies and reduced reciprocation in consecutive grooming bouts within a session (Surbeck & Hohmann, 2015). Put differently, if a relationship is already valuable there may be less pressure to restore balance instantly. On the contrary, valuable partners could reciprocate more quickly to maintain a preferred relationship. Evidence exists where grooming given and received are time-matched within bouts when both individuals provided grooming (Chancellor & Isbell, 2009; Barrett et al., 2002, 1999). Theoretically, both could be possible. Prior work has also now repeatedly shown that grooming exchanges do not just occur in strict bout-to-bout symmetry and that reciprocation extends over longer time windows across primate taxa (Frank & Silk, 2009; Gomes et al., 2009; Manson et al., 2004). Equally, the emotional mediation framework predicts that emotional attachments to certain partners shape how individuals respond to cooperative interactions (Aureli & Schaffner, 2002). Closer bonds can be buffered by trust and familiarity and could plausibly absorb temporary imbalance without threatening the bond, expressing reciprocity more loosely in time (Jaeggi et al., 2013; Seyfarth & Cheney, 2012; Schino & Aureli, 2009). Our key point is that the relationship is still reciprocal, but the mechanism of how reciprocity is maintained appears to be faster return to balance between strong bonds rather than prolonged tolerance of imbalance in crested macaques.

To reinforce our interpretation, substantial variance was attributed to dyad and individual identity, which suggests the timing of reciprocation is individual and relationship specific. Our results were not simply determined by bond strength alone, but also by pair-specific traits, and to a lesser degree individual characteristics, that made some dyads repeatedly reciprocate more quickly or more slowly. We did have to exclude dominance rank, relatedness, sex, and age as fixed effects, a common limitation of captive studies with small sample sizes, although it is likely that these factors also contributed to variation in grooming reciprocity. Some of this influence may have been captured in our random effects but we state this with some uncertainty. However, these factors are known to affect grooming and social behaviour more generally (Schino & Aureli, 2008a; Cooper & Bernstein, 2000). A new breeding male had also been recently introduced to the CERZ group which could have unsettled social dynamics. Similarly, meta-analyses assessing individual personality differences across taxa have shown the evolutionary significance of risky or explorative behaviours that could help overall fitness and subsequently affect social relationships (Moiron et al., 2019; Smith & Blumstein, 2008). What we can more clearly state is that some of the variance could be attributed to sex differences. CERZ females reciprocated significantly faster than males. This suggests sex-specific social strategies may shape the temporal structure of grooming reciprocation. For example, Assamese macaque (*M. assamensis)* females groomed each other more than male-male dyads and were involved in more simultaneous multi-partner grooming interactions than males (Cooper & Bernstein, 2000). Female primates are known to employ grooming more strategically than males to shape and maintain relationship quality (Roubová et al., 2015; Silk et al., 2013). Combined with evidence that female macaques show greater inhibitory control and are less easily distracted during cognitive tasks than males (Loyant et al., 2021), this suggests that sex differences in both social strategy and cognitive tendency may together contribute to the faster grooming reciprocation observed in CERZ females. Whilst we do not claim a causal link, these findings are consistent with the possibility that sex-specific grooming strategies and impulse control are related in a way that affects how quickly females reciprocate and restore grooming balance. Sex differences in temporal grooming structures would be beneficial to understand in the sense that faster reciprocation may not just reflect relationship quality, but also individual differences in social responsiveness. This remains a tentative explanation given that captive groups experience different social pressures, reduced ranging opportunities, and are assembled by humans rather than forming naturally; testing this in wild populations would be a necessary next step.

Grooming reciprocity may also not always be a grooming-for-grooming type exchange. Primate exchanges have linked preferential grooming to other social commodities such as tolerance, sex, coalitionary support, and reduced aggression (Kaburu & Newton-Fisher, 2015; Molesti & Majolo, 2015; Tiddi et al., 2011; Frank & Silk, 2009; Gumert, 2007; Schino et al., 2007; Barrett & Henzi, 2006; de Waal, 1997). We cannot exclude the possibility that some of the reciprocity we observed was in part a reflection of multi-commodity exchanges during a specific period of time. Previous macaque studies have found evidence of grooming being exchanged for other social commodities, e.g., coalitionary support or cofeeding (Molesti & Majolo, 2015; Schino et al., 2007; Ventura et al., 2006). One possible approach would be to leverage the controlled conditions of captive environments to manipulate contexts in which grooming can be exchanged for other commodities, and to observe the time frames over which exchanges occur. For example, Hemelrijk (1994) induced food-based conflict in a captive group of long-tailed macaques and tested whether being groomed increased the likelihood of receiving agonistic support. Monkeys were significantly more likely to support recent groomers. This would be useful to investigate to understand if a wider social economy contributes to quicker reciprocation latencies rather than narrow, grooming-only exchanges.

At the same time, dyads may show an imbalance across consecutive grooming bouts whilst remaining reciprocally balanced over a longer time window, a distinction that has implications for how results could be interpreted. We observed a large proportion of unilateral grooming episodes between reciprocal events, i.e., monkey A groomed monkey B on multiple occasions before monkey B reciprocated. In another study with crested macaques, grooming episodes were not time-matched (Tyrrell & Berman, 2012). Whether balance instead emerges across a longer time frame remains unknown and a question for the future. If bond strength shapes the time frame over which reciprocity occurs as we observed here, strongly bonded dyads may tolerate greater short-term imbalances because the relationship provides greater longer-term assurance. This makes the within-and-across-bout structure of reciprocity a theoretically important question, not solely a methodological one. Evidence for both grooming being time-matched in shorter-time frames and grooming becoming more balanced over consecutive bouts exists across primate species (Surbeck & Hohmann, 2015; Frank & Silk, 2009; Payne et al., 2003; Barrett et al., 2002, 1999). This variance is attributed largely to species differences in socioecology or demographic factors (Sánchez-Amaro & Amici, 2015). This is why the question of time frame is so important to distinguish and has been highlighted as an important baseline to establish prior to testing conditions of a primate biological market (Dunayer et al., 2019; Sánchez-Amaro & Amici, 2015). We did not test whether social bond strength predicted reciprocation differently within versus across bouts because we did not assign bout membership for individual grooming episodes. But we did address the initial question of assessing reciprocation over both the short and long term and its social bond effects rather than narrower analysis of more immediate bout-level definitions. It would be interesting to assess the temporal dynamics of shorter-term imbalances coexisting with our results of stronger bonds reciprocating faster. A dyad temporarily uneven at the bout level may show a rapid tendency to create balance particularly when partners are more strongly bonded. That would temporarily permit imbalance but then ensure a return to balance if the relationship is of strong value. A within and consecutive bout perspective to rebalancing grooming asymmetry may complement our approach in future studies.

### Eigenvector centrality effects on reciprocation

We also investigated whether individuals’ broader EC contributed to explaining the observed variance in reciprocation latency. EC weights an individual’s connections by the centrality of their partners, meaning individuals connected to other well-connected individuals score higher (Brent, 2015). Upon analysis, we found that degree carried no discriminatory value because both networks were fully connected: every individual was connected to every other group member. Furthermore, strength correlated closely with EC. We chose EC over strength because strength duplicates what our CSI already captures at the dyad level, whereas EC indexes something our CSI did not. Under BMT, the rate at which grooming is exchanged depends on the alternatives available to both partners rather than on the dyad in isolation (Noë & Hammerstein, 1994). Two individuals with equivalent grooming investment differ if one grooms with well-connected partners whose grooming is in demand elsewhere, and the other with peripheral partners. If partner competition shapes exchange rates, that difference should be visible in reciprocation timing. Contrary to our initial hypothesis, we did not observe a credible effect of EC on reciprocation latency. LOO cross validation indicated that EC did not lead to predictive improvements compared to a model alone including only the social bond strength as a predictor. These results suggest that reciprocal timing is more likely determined by the localised strength of the dyadic relationship than by the connectedness of the partners to whom an individual is indirectly linked. This aligns with the consensus that grooming reciprocity is better understood as a property of relationship specific dynamics. EC may still matter indirectly, but not after bond strength was accounted for.

The lack of influence of EC on grooming reciprocity may reflect the tolerance style in crested macaques. In more despotic macaque species (grade 1 - 2), social systems are more constrained by dominance, with greater affiliation concentrated around higher ranking individuals and clustered within kin groups. In these systems, network position is more heavily dictated by rank: top-ranking individuals tend to be more central compared to other group members (Wooddell et al., 2020; Sueur et al., 2011). However, this rank-centrality link is not universal when it comes to grooming: middle ranking Nicobar long-tailed macaques *(M. fascicularis umbrosus)* were the major contributors to the group’s grooming distribution (Mishra et al., 2020). Yet, in tolerant systems (grade 3 – 4), social structures are less constrained by nepotism or steep hierarchical arrangements, and greater degrees of freedom are observed in crested macaques’ interactions and relationships (Duboscq et al., 2013; Thierry, 2007). It is possible that EC may be a less sensitive measure for assessing temporal grooming patterns in more tolerant macaques if the social distributions of their networks are more freely arranged. Differences in cognition are present across macaque tolerance gradients. Cooperation is jointly shaped by prosocial tendencies, kinship, dyadic tolerance, and individual gregariousness (Bhattacharjee et al., 2026; Neumann & Fischer, 2026; Brosnan & de Waal, 2002). Yet cooperative interactions in despotic systems are more concentrated around a subset of strongly bonded individuals with interdependent relationships (Bhattacharjee et al., 2026). More tolerant macaque species similarly outperform less tolerant ones in social cognition and inhibitory control tasks (Joly et al., 2017). Whilst EC may be a good indicator for broader individual social embeddedness at group level, it may be less influential in describing proximate mechanisms of grooming dynamics in more tolerant social systems. Temporal grooming strategies seem more likely to be dictated by the maintenance of partner-specific investment described by BMT (Noë & Hammerstein, 1994). However, our results are limited by the number of observed individuals. This makes it difficult to determine if the effects here are a biological feature of tolerant macaque species or more likely a sampling limitation. We could only use EC to assess centrality effects. Degree carried no variance in either group, and strength correlated with EC at r = 0.95 in CERZ and r = 0.97 in EMER. The restricted range of EC scores is also reflected through the wide credible intervals. In captivity, restricted space forces individuals closer together. The resulting networks can then reflect who was available rather than who was preferred (De Moor et al., 2025). This lack of variation in individual EC ultimately was a contributory factor towards masking any potential temporal-latency effects of centrality (De Moor et al., 2025). It may be of even more significance to ensure a greater number of comparative groups and centrality scores in tolerant macaque species to more rigorously test for genuine effects, given their more egalitarian social structure. It is possible that some centrality measures could more reliably predict ability in other cognitive domains than the social and dyadic measures tested here. Whilst eigenvector centrality did not help to predict temporal reciprocity durations, this remains to be fully tested. This would all require a much more robust and comparative data set than ours, with the inclusion of factors we were unable to measure such as dominance ranking, kinship or age.

## Conclusion

To summarise, using survival analyses we showed that in two captive groups of crested macaques, most reciprocation occurred over longer timescales than commonly assumed. Stronger bonds predicted faster reciprocation, indicating that the quality of the dyadic relationship is a key driver for temporal reciprocation windows. These findings support a biological markets interpretation and highlight the behavioural significance of including unilateral grooming episodes in analyses as biologically relevant social interactions. Grooming reciprocity should not be assumed over shorter-term windows only, as relevant time scales of when they become balanced and reciprocal may vary with species’ social style and dyadic factors. Future research should focus on testing temporal patterns in wild populations where social pressures and group sizes are more variable. Investigating whether latency is better predicted by other dyadic variables such as rank distance and the exchange of other social commodities across the macaque tolerance gradient.

## Funding

This research was supported by internal funding via a Faculty of Sciences and Health PhD bursary from the University of Portsmouth.

## Acknowledgements

We thank Zoo de CERZA and Emerald Park for allowing us to conduct this research and for their support throughout data collection.

## Notes

https://github.com/lukecollins06/buildingblocksofcooperation-THESIS

### Competing Interest Statement

The authors have declared no competing interest.

### Summary of Updates

The version of the manuscript has updated authors, fixed citations, and some grammar corrections.

