## Supplementary Tables & Figures for "Temporal dynamics of grooming reciprocity in crested macaques: Strongly bonded individuals reciprocate grooming more quickly"

### Supplementary Table S1: full summary of social bond strength (CSI) scores for CERZ and EMER

#### CERZ

| focal_id1 | focal_id2 | dyad | CSI_affil | CSI_groom | CSI_prox | CSI_total | z_CSI |
|---|---|---|---|---|---|---|---|
| BROW | BUNTA | BROW_BUNTA | 0.3355111 | 0 | 0 | 0.11183704 | -0.97127612 |
| BROW | FIDGET | BROW_FIDGET | 0 | 0.52354344 | 0.2184856 | 0.24734301 | -0.80233961 |
| BROW | KAOS | BROW_KAOS | 0.6452137 | 2.76537567 | 0.1848724 | 1.19848726 | 0.38346046 |
| BROW | MAKASSAR | BROW_MAKASSAR | 0.1747454 | 1.26728671 | 0.2002785 | 0.54743685 | -0.42820993 |
| BROW | MALI | BROW_MALI | 0 | 1.27383753 | 0.4450632 | 0.57296692 | -0.39638136 |
| BROW | MONK | BROW_MONK | 1.1852295 | 0.16296823 | 1.5673966 | 0.97186477 | 0.10092818 |
| BROW | QUANNY | BROW_QUANNY | 0 | 0.03913023 | 0.6729356 | 0.23735528 | -0.81479141 |
| BROW | SULA | BROW_SULA | 0.1352867 | 0 | 0 | 0.04509558 | -1.0544833 |
| BROW | SUNDA | BROW_SUNDA | 0.8065171 | 0.7074318 | 0.8319259 | 0.78195827 | -0.13582997 |
| BROW | TALIA | BROW_TALIA | 0.9174132 | 0.15320062 | 0.6759398 | 0.58218454 | -0.38488967 |
| BUNTA | FIDGET | BUNTA_FIDGET | 0.911715 | 1.17967282 | 0.7314517 | 0.94094652 | 0.06238211 |
| BUNTA | KAOS | BUNTA_KAOS | 0.6213169 | 0.73048508 | 0.8901265 | 0.74730948 | -0.17902693 |
| BUNTA | MAKASSAR | BUNTA_MAKASSAR | 0.2516333 | 0 | 0.288401 | 0.18001144 | -0.88628249 |
| BUNTA | MALI | BUNTA_MALI | 0.4493452 | 1.05374374 | 0.7725026 | 0.75853053 | -0.16503754 |
| BUNTA | MONK | BUNTA_MONK | 0.6989815 | 0.30391456 | 0.7009746 | 0.56795688 | -0.40262742 |
| BUNTA | QUANNY | BUNTA_QUANNY | 2.1775962 | 1.21668887 | 3.7898846 | 2.39472322 | 1.87481858 |
| BUNTA | SULA | BUNTA_SULA | 6.6840105 | 3.27356717 | 2.4784459 | 4.14534119 | 4.05732972 |
| BUNTA | SUNDA | BUNTA_SUNDA | 1.8639506 | 0.57712943 | 2.4923541 | 1.64447806 | 0.9394812 |
| BUNTA | TALIA | BUNTA_TALIA | 2.4146633 | 1.70236598 | 1.0195994 | 1.71220957 | 1.02392269 |
| FIDGET | KAOS | FIDGET_KAOS | 1.397963 | 2.82889081 | 1.9026453 | 2.04316637 | 1.4365295 |
| FIDGET | MAKASSAR | FIDGET_MAKASSAR | 0.4765783 | 0.27136657 | 0.546214 | 0.43138628 | -0.57289122 |
| FIDGET | MALI | FIDGET_MALI | 2.8518445 | 2.8133659 | 0.1922673 | 1.95249257 | 1.32348565 |
| FIDGET | MONK | FIDGET_MONK | 0.399418 | 0.15489866 | 0.4577793 | 0.33736533 | -0.69010798 |
| FIDGET | QUANNY | FIDGET_QUANNY | 1.1852295 | 0.92194166 | 1.0449311 | 1.05070074 | 0.19921368 |
| FIDGET | SULA | FIDGET_SULA | 1.3015517 | 0.47666653 | 1.07736 | 0.95185941 | 0.07598731 |
| FIDGET | SUNDA | FIDGET_SUNDA | 0 | 0.14690839 | 0.7009746 | 0.28262766 | -0.75834992 |
| FIDGET | TALIA | FIDGET_TALIA | 1.3280648 | 1.06755312 | 1.6022276 | 1.3326152 | 0.55067897 |
| KAOS | MAKASSAR | KAOS_MAKASSAR | 0.1613034 | 0 | 0 | 0.05376781 | -1.04367156 |
| KAOS | MALI | KAOS_MALI | 0.8677012 | 0.48227237 | 0.8287384 | 0.72623732 | -0.20529777 |
| KAOS | MONK | KAOS_MONK | 0 | 1.03387099 | 0.5768019 | 0.53689098 | -0.44135756 |
| KAOS | QUANNY | KAOS_QUANNY | 0.3106584 | 0 | 0.7121012 | 0.34091987 | -0.6856765 |
| KAOS | SULA | KAOS_SULA | 0.6989815 | 0.11723134 | 0.5097997 | 0.44200418 | -0.55965378 |
| KAOS | SUNDA | KAOS_SUNDA | 0.3744544 | 0.32632139 | 0.4291681 | 0.37664796 | -0.64113397 |
| KAOS | TALIA | KAOS_TALIA | 0.3083742 | 0.14775466 | 1.6964763 | 0.71753506 | -0.21614696 |
| MAKASSAR | MALI | MAKASSAR_MALI | 0.5436523 | 1.79630631 | 0.2670379 | 0.86899884 | -0.02731571 |

| | | | | | | | |
|---|---|---|---|---|---|---|---|
| MAKASSAR | MONK | MAKASSAR_MONK | 0.182343 | 0.02469797 | 0.3134793 | 0.17350676 | -0.89439193 |
| MAKASSAR | QUANNY | MAKASSAR_QUANNY | 0 | 0.3276377 | 0.288401 | 0.20534622 | -0.85469738 |
| MAKASSAR | SULA | MAKASSAR_SULA | 0.1352867 | 0.04656616 | 0.1550543 | 0.1123024 | -0.97069596 |
| MAKASSAR | SUNDA | MAKASSAR_SUNDA | 0.3226068 | 0.40160904 | 0.8319259 | 0.51871392 | -0.46401906 |
| MAKASSAR | TALIA | MAKASSAR_TALIA | 1.1140018 | 2.56801969 | 0.3004177 | 1.32747971 | 0.54427651 |
| MALI | MONK | MALI_MONK | 0.3226068 | 0.04968021 | 0.3697448 | 0.24734397 | -0.80233842 |
| MALI | QUANNY | MALI_QUANNY | 0.4493452 | 0 | 0 | 0.14978175 | -0.92397011 |
| MALI | SULA | MALI_SULA | 1.5418709 | 0.05366378 | 0.2827461 | 0.62609359 | -0.33014786 |
| MALI | SUNDA | MALI_SUNDA | 1.3738602 | 0.06286193 | 0.1657477 | 0.5341566 | -0.44476654 |
| MALI | TALIA | MALI_TALIA | 2.9357223 | 0.31091453 | 1.1673373 | 1.47132469 | 0.72360934 |
| MONK | QUANNY | MONK_QUANNY | 0 | 0.63354232 | 0.7009746 | 0.44483897 | -0.55611962 |
| MONK | SULA | MONK_SULA | 0.3494907 | 0.87381848 | 0.1602228 | 0.46117733 | -0.53575044 |
| MONK | SUNDA | MONK_SUNDA | 0.6710222 | 0.1745001 | 0.7690693 | 0.5381972 | -0.43972908 |
| MONK | TALIA | MONK_TALIA | 0.541147 | 4.63247945 | 1.4730157 | 2.21554738 | 1.65143845 |
| QUANNY | SULA | QUANNY_SULA | 0.9829427 | 1.77463683 | 3.8303254 | 2.19596833 | 1.62702908 |
| QUANNY | SUNDA | QUANNY_SUNDA | 0 | 0.21012636 | 2.8484047 | 1.01951035 | 0.16032834 |
| QUANNY | TALIA | QUANNY_TALIA | 0.6354377 | 0.6002857 | 1.383742 | 0.87315516 | -0.02213399 |
| SULA | SUNDA | SULA_SUNDA | 1.0167004 | 5.20799729 | 2.9131412 | 3.04594627 | 2.6867042 |
| SULA | TALIA | SULA_TALIA | 1.0215883 | 0.275133 | 0.9859862 | 0.76090252 | -0.16208035 |
| SUNDA | TALIA | SUNDA_TALIA | 0.9251226 | 1.25613888 | 1.5551033 | 1.24545491 | 0.44201546 |

### EMER

| focal_id1 | focal_id2 | dyad | CSI_affil | CSI_groom | CSI_prox | CSI_total | z_CSI |
|---|---|---|---|---|---|---|---|
| BASUKI | BUMI | BASUKI_BUMI | 3.416533 | 1.45995311 | 1.6109224 | 2.1624695 | 2.10154478 |
| BASUKI | DOUGIE | BASUKI_DOUGIE | 0.76276086 | 0 | 0.9582923 | 0.5736844 | -0.63969858 |
| BASUKI | DRUSILLA | BASUKI_DRUSILLA | 0.07478048 | 0 | 0.2332557 | 0.1026787 | -1.45235793 |
| BASUKI | EKAH | BASUKI_EKAH | 0.63563405 | 0.12971037 | 0.7744819 | 0.5132754 | -0.74392639 |
| BASUKI | INDAH | BASUKI_INDAH | 0.57412107 | 0.23595075 | 0.6395721 | 0.4832147 | -0.79579238 |
| BASUKI | KERANA | BASUKI_KERANA | 2.05043241 | 0 | 0.7035294 | 0.9179873 | -0.04564846 |
| BASUKI | MASAMBA | BASUKI_MASAMBA | 0.55617979 | 1.20711947 | 1.9207151 | 1.2280048 | 0.48924673 |
| BASUKI | SETANA | BASUKI_SETANA | 0.24605189 | 1.11598078 | 1.2151871 | 0.8590733 | -0.14729696 |
| BUMI | DOUGIE | BUMI_DOUGIE | 0.7150883 | 0.48123497 | 0.2168549 | 0.4710594 | -0.8167647 |
| BUMI | DRUSILLA | BUMI_DRUSILLA | 0.35313003 | 0.09877696 | 0.1927599 | 0.214889 | -1.25875367 |
| BUMI | EKAH | BUMI_EKAH | 0.89736571 | 0.28086725 | 0.6997672 | 0.626 | -0.54943465 |
| BUMI | INDAH | BUMI_INDAH | 0.23113965 | 0.43701442 | 0.5407292 | 0.4029611 | -0.93425955 |
| BUMI | KERANA | BUMI_KERANA | 1.23274482 | 0.06044007 | 0.3304456 | 0.5412102 | -0.69572863 |

| | | | | | | | |
|---|---|---|---|---|---|---|---|
| BUMI | MASAMBA | BUMI_MASAMBA | 0 | 0.37048149 | 1.8368888 | 0.7357901 | -0.36000609 |
| BUMI | SETANA | BUMI_SETANA | 0.61637241 | 2.00979234 | 1.9225926 | 1.5162525 | 0.98658083 |
| DOUGIE | DRUSILLA | DOUGIE_DRUSILLA | 1.19648762 | 4.26946498 | 3.0031675 | 2.82304 | 3.24127385 |
| DOUGIE | EKAH | DOUGIE_EKAH | 1.43017661 | 1.43294263 | 0.9913368 | 1.2848187 | 0.58727177 |
| DOUGIE | INDAH | DOUGIE_INDAH | 0.98420756 | 2.29622865 | 0.9273796 | 1.4026053 | 0.79049729 |
| DOUGIE | KERANA | DOUGIE_KERANA | 0.90219026 | 0.05443443 | 0.7994652 | 0.5853633 | -0.61954812 |
| DOUGIE | MASAMBA | DOUGIE_MASAMBA | 0.31781702 | 0 | 1.0842747 | 0.4673639 | -0.82314083 |
| DOUGIE | SETANA | DOUGIE_SETANA | 0.98420756 | 1.48498414 | 0.7674866 | 1.0788928 | 0.23197319 |
| DRUSILLA | EKAH | DRUSILLA_EKAH | 0.70626005 | 2.89226632 | 1.3768567 | 1.658461 | 1.23194332 |
| DRUSILLA | INDAH | DRUSILLA_INDAH | 0.43586335 | 1.98987615 | 0.8497173 | 1.0918189 | 0.25427562 |
| DRUSILLA | KERANA | DRUSILLA_KERANA | 0.36321946 | 1.36930847 | 0.9630129 | 0.8985136 | -0.07924771 |
| DRUSILLA | MASAMBA | DRUSILLA_MASAMBA | 0 | 0.06091199 | 0.826114 | 0.2956753 | -1.11936724 |
| DRUSILLA | SETANA | DRUSILLA_SETANA | 0.65379502 | 1.35352524 | 0.5664782 | 0.8579328 | -0.14926463 |
| EKAH | INDAH | EKAH_INDAH | 1.38683792 | 0.112537 | 0.5106887 | 0.6700212 | -0.47348184 |
| EKAH | KERANA | EKAH_KERANA | 3.1589086 | 0.20153318 | 1.411904 | 1.5907819 | 1.11517177 |
| EKAH | MASAMBA | EKAH_MASAMBA | 1.12170714 | 0.19242838 | 1.5744761 | 0.9628706 | 0.03179187 |
| EKAH | SETANA | EKAH_SETANA | 1.23274482 | 0.56373765 | 0.9612963 | 0.9192596 | -0.0434532 |
| INDAH | KERANA | INDAH_KERANA | 1.66853937 | 1.64992558 | 0.5886062 | 1.3023571 | 0.61753197 |
| INDAH | MASAMBA | INDAH_MASAMBA | 0.53932586 | 0.15843007 | 0.6008102 | 0.4328554 | -0.88268081 |
| INDAH | SETANA | INDAH_SETANA | 2.46308193 | 1.71113456 | 1.0842747 | 1.7528304 | 1.39476545 |
| KERANA | MASAMBA | KERANA_MASAMBA | 1.46388447 | 1.14966705 | 0.9612963 | 1.1916159 | 0.42646248 |
| KERANA | SETANA | KERANA_SETANA | 0.39727128 | 0.04104059 | 0.4027306 | 0.2803475 | -1.14581347 |
| MASAMBA | SETANA | MASAMBA_SETANA | 0.23113965 | 1.12830094 | 1.9526332 | 1.1040246 | 0.2753349 |

**Supplementary Table S2: Tweedie GLMM results: effect of dyadic grooming rate received on grooming rate given**

Family: tweedie ( log )

Formula: given_rate ~ received_rate + Group + (1 | dyad_id)

Data: groom_paired_combined

| AIC | BIC | logLik | -2*log(L) | df.resid |
|---|---|---|---|---|
| 714.3 | 729.0 | -351.1 | 702.3 | 80 |

Random effects:

Conditional model:

| Groups | Name | Variance | Std.Dev. |
|---|---|---|---|
| dyad_id | (Intercept) | 3.097e-08 | 0.000176 |

Number of obs: 86, groups: dyad_id, 86

Dispersion parameter for tweedie family (): 2.84

Conditional model:

<table>
<tr><th></th><th>Estimate</th><th>Std. Error</th><th>z value</th><th>Pr(>|z|)</th><th></th></tr>
<tr><td>(Intercept)</td><td>2.270169</td><td>0.183800</td><td>12.351</td><td>< 2e-16</td><td>***</td></tr>
<tr><td>received_rate</td><td>0.017260</td><td>0.002909</td><td>5.933</td><td>2.98e-09</td><td>***</td></tr>
<tr><td>GroupEMER</td><td>0.743450</td><td>0.223567</td><td>3.325</td><td>0.000883</td><td>***</td></tr>
</table>

---

Signif. codes: 0 ‘***’ 0.001 ‘**’ 0.01 ‘*’ 0.05 ‘.’ 0.1 ‘ ’ 1

**Supplementary Figure S1: Kaplan-Meier curves of time to grooming reciprocation by sex for CERZ and EMER**

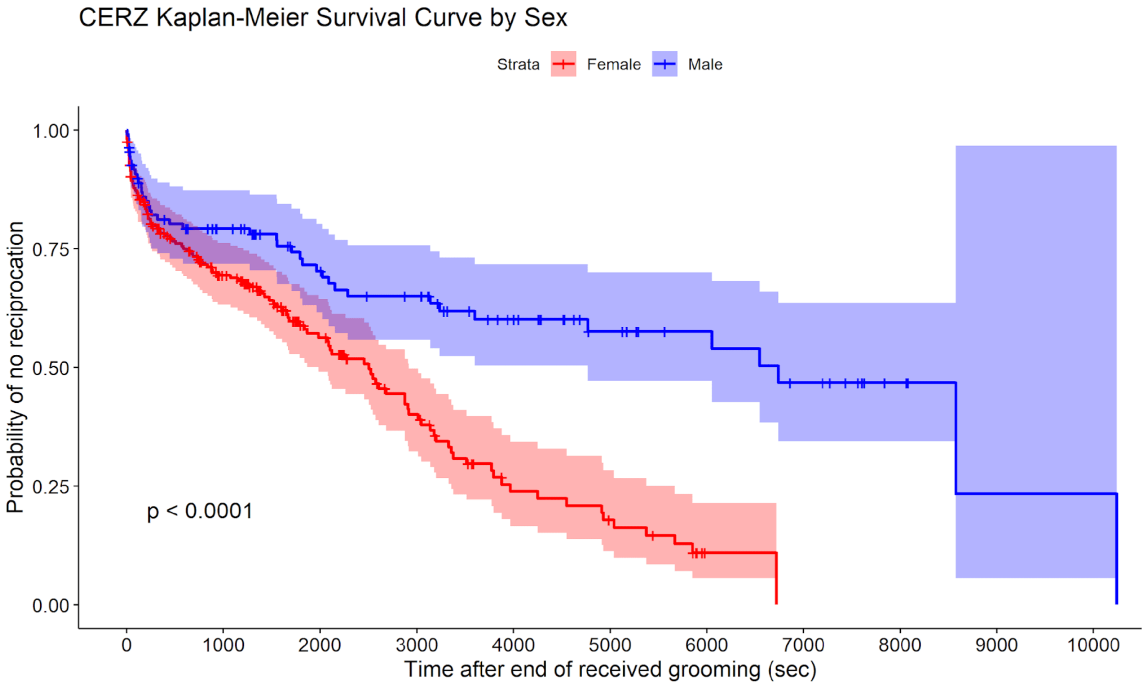


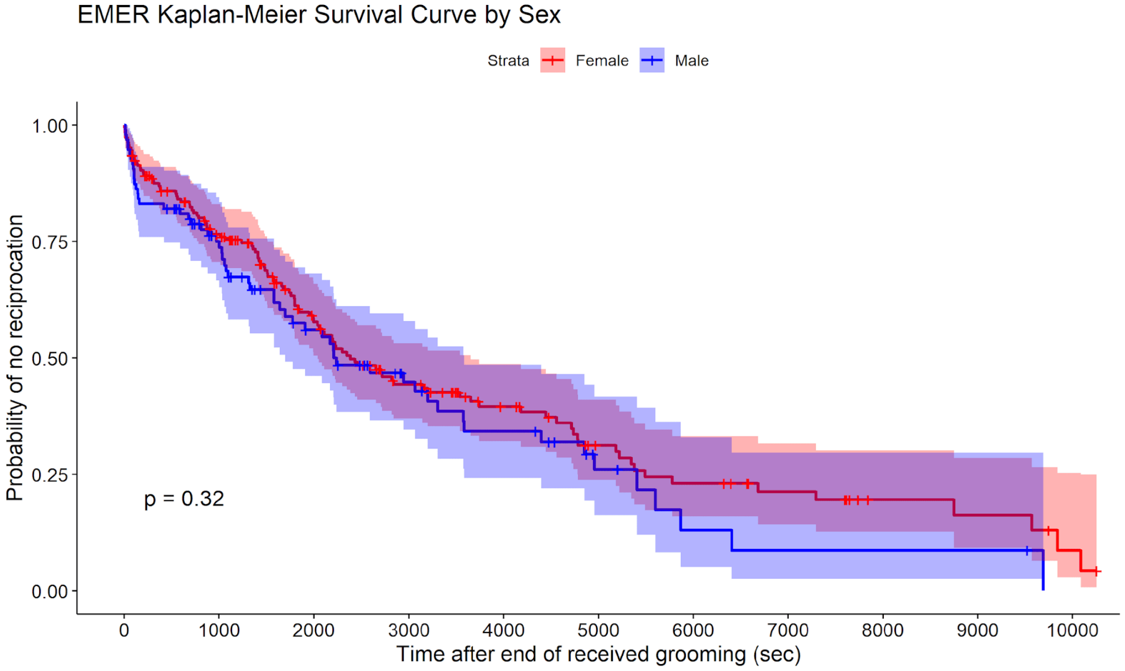

**Supplementary Table S3: Bayesian gamma survival model results: effect of social bond strength on reciprocation latency**

Family: gamma
Links: mu = log; shape = identity
Formula: time | cens(censored) ~ z_CSI + (1 | mm(focal_id, partner_id)) + (1 | dyad_index) + (1 | group_ID)
Data: CERZ_EMER_grooming_data_combined (Number of observations: 597)
Draws: 4 chains, each with iter = 4000; warmup = 2000; thin = 1;
total post-warmup draws = 8000

Multilevel Hyperparameters:

**~dyad_index** (Number of levels: 86)

| | Estimate | Est.Error | l-95% CI | u-95% CI | Rhat | Bulk_ESS | Tail_ESS |
|---|---|---|---|---|---|---|---|
| sd(Intercept) | **0.96** | 0.16 | **0.67** | **1.31** | 1.00 | 2356 | 4066 |

**~group_ID (**Number of levels: 2)

| | Estimate | Est.Error | l-95% CI | u-95% CI | Rhat | Bulk_ESS | Tail_ESS |
|---|---|---|---|---|---|---|---|
| sd(Intercept) | **2.58** | 1.20 | **1.15** | **5.57** | 1.00 | 4314 | 4849 |

**~mmfocal_idpartner_id** (Number of levels: 20)

| | Estimate | Est.Error | l-95% CI | u-95% CI | Rhat | Bulk_ESS | Tail_ESS |
|---|---|---|---|---|---|---|---|
| sd(Intercept) | **0.56** | 0.35 | **0.03** | **1.30** | 1.00 | 1189 | 3073 |

Regression Coefficients:

| | Estimate | Est.Error | l-95% CI | u-95% CI | Rhat | Bulk_ESS | Tail_ESS |
|---|---|---|---|---|---|---|---|
| Intercept | 5.79 | 0.51 | 4.78 | 6.81 | 1.00 | 10997 | 6016 |
| **z_CSI** | **-0.38** | 0.14 | **-0.67** | **-0.11** | 1.00 | 4226 | 4668 |

Further Distributional Parameters:

| | Estimate | Est.Error | l-95% CI | u-95% CI | Rhat | Bulk_ESS | Tail_ESS |
|---|---|---|---|---|---|---|---|
| shape | 0.69 | 0.04 | 0.61 | 0.78 | 1.00 | 14669 | 5372 |

Draws were sampled using sampling(NUTS). For each parameter, Bulk_ESS and Tail_ESS are effective sample size measures, and Rhat is the potential scale reduction factor on split chains (at convergence, Rhat = 1).

**Supplementary Figure S2: posterior predictive checks (density overlay) of observed versus replicated reciprocation latencies for the gamma (top) and hurdle lognormal (bottom) models**

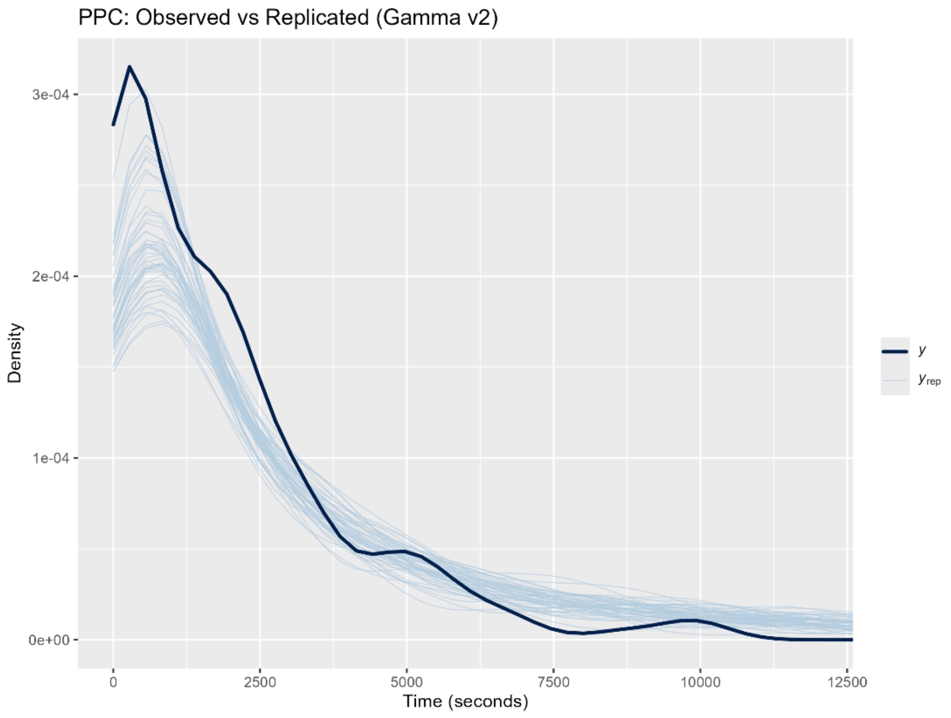


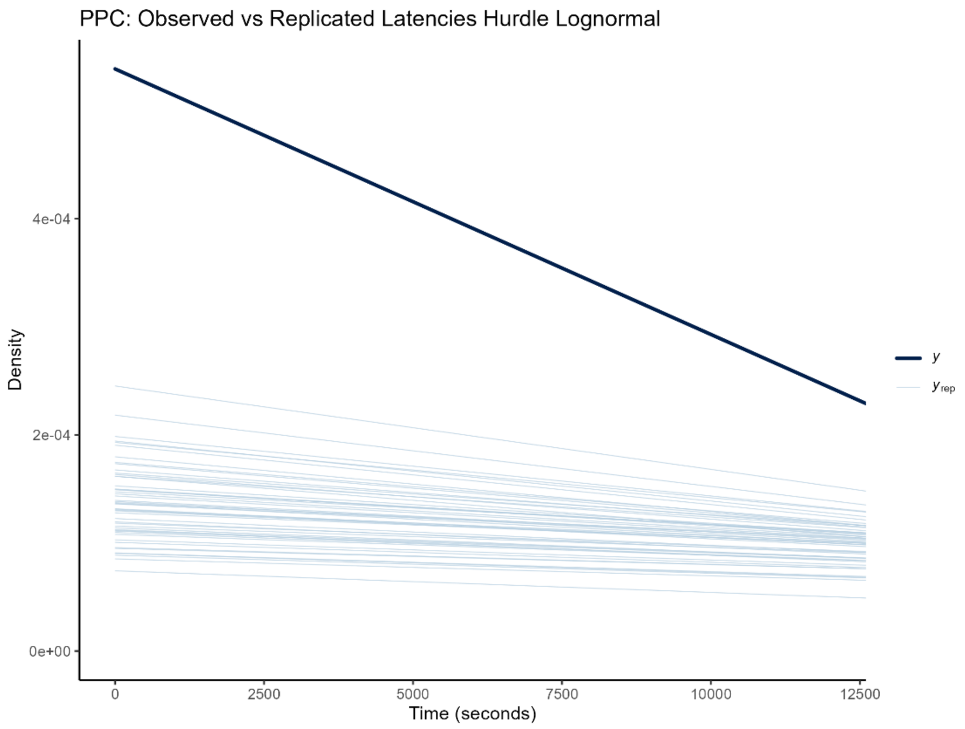

**Supplementary Table S4: degree, strength, and eigenvector centrality scores for CERZ and EMER**

**CERZ**

| individual | degree | strength | eigenvector |
|---|---|---|---|
| TALIA | 10 | 11.59458 | 0.751782083 |
| SULA | 10 | 11.4058 | 1 |
| BUNTA | 10 | 10.96405 | 0.965857751 |
| SUNDA | 10 | 8.635582 | 0.747007413 |
| FIDGET | 10 | 8.378279 | 0.600315734 |
| QUANNY | 10 | 8.311783 | 0.75044501 |
| MALI | 10 | 6.356186 | 0.442551202 |
| KAOS | 10 | 5.738631 | 0.395831118 |
| MONK | 10 | 5.640825 | 0.428282189 |
| BROW | 10 | 4.423302 | 0.274280271 |
| MAKASSAR | 10 | 2.878101 | 0.200782844 |

**EMER**

| individual | degree | strength | eigenvector |
|---|---|---|---|
| DOUGIE | 8 | 12.1396 | 1 |
| BASUKI | 8 | 9.236691 | 0.771037 |
| EKAH | 8 | 8.484125 | 0.737867 |
| BUMI | 8 | 8.271725 | 0.69751 |
| DRUSILLA | 8 | 8.245629 | 0.781551 |
| MASAMBA | 8 | 8.106715 | 0.679775 |
| SETANA | 8 | 8.064853 | 0.687224 |
| KERANA | 8 | 7.581289 | 0.645801 |
| INDAH | 8 | 6.923958 | 0.613511 |

**Supplementary Table S5: Bayesian gamma survival model results: effects of social bond strength and eigenvector centrality on reciprocation latency**

Family: gamma
Links: mu = log; shape = identity
Formula: time | cens(censored) ~ z_CSI + z_eigenvector + (1 | mm(focal_id, partner_id)) + (1 | dyad_index) + (1 | group_ID)
Data: CERZ_EMER_centrality_groom_combined (Number of observations: 597)
Draws: 4 chains, each with iter = 4000; warmup = 2000; thin = 1;
total post-warmup draws = 8000

Multilevel Hyperparameters:

**~dyad_index** (Number of levels: 86)

| | Estimate | Est.Error | l-95% CI | u-95% CI | Rhat | Bulk_ESS | Tail_ESS |
|---|---|---|---|---|---|---|---|
| sd(Intercept) | **0.98** | 0.17 | **0.67** | **1.33** | 1.00 | 2264 | 3873 |

**~group_ID** (Number of levels: 2)

| | Estimate | Est.Error | l-95% CI | u-95% CI | Rhat | Bulk_ESS | Tail_ESS |
|---|---|---|---|---|---|---|---|
| sd(Intercept) | **2.58** | 1.25 | **1.12** | **5.66** | 1.00 | 3715 | 5033 |

**~mmfocal_idpartner_id** (Number of levels: 20)

| | Estimate | Est.Error | l-95% CI | u-95% CI | Rhat | Bulk_ESS | Tail_ESS |
|---|---|---|---|---|---|---|---|
| sd(Intercept) | **0.58** | 0.36 | **0.03** | **1.35** | 1.00 | 1001 | 2572 |

Regression Coefficients:

| | Estimate | Est.Error | l-95% CI | u-95% CI | Rhat | Bulk_ESS | Tail_ESS |
|---|---|---|---|---|---|---|---|
| Intercept | 5.80 | 0.53 | 4.78 | 6.84 | 1.00 | 9753 | 5964 |
| **z_CSI** | **-0.40** | 0.15 | **-0.69** | **-0.12** | 1.00 | 3954 | 5173 |
| **z_eigenvector** | **0.05** | 0.16 | **-0.25** | **0.36** | 1.00 | 5446 | 5415 |

Further Distributional Parameters:

| | Estimate | Est.Error | l-95% CI | u-95% CI | Rhat | Bulk_ESS | Tail_ESS |
|---|---|---|---|---|---|---|---|
| shape | 0.69 | 0.04 | 0.61 | 0.77 | 1.00 | 12674 | 5978 |

Draws were sampled using sampling(NUTS). For each parameter, Bulk_ESS and Tail_ESS are effective sample size measures, and Rhat is the potential scale reduction factor on split chains (at convergence, Rhat = 1).

**Supplementary Table S6: LOO comparison of gamma models with and without eigenvector centrality**

Moment matching was applied because one of 597 observations exceeded the pareto <k threshold of 0.7; this did not affect predictions.

| | elpd_diff | se_diff |
|---|---|---|
| CERZ_EMER_brm_gamma_v2 | 0.0 | 0.0 |
| CERZ_EMER_brm_gamma_eigenvector | -0.1 | 0.3 |

**Supplementary Figure S3: posterior predictive check (density overlay) of observed versus replicated reciprocation latencies for the social bond strength and eigenvector model**

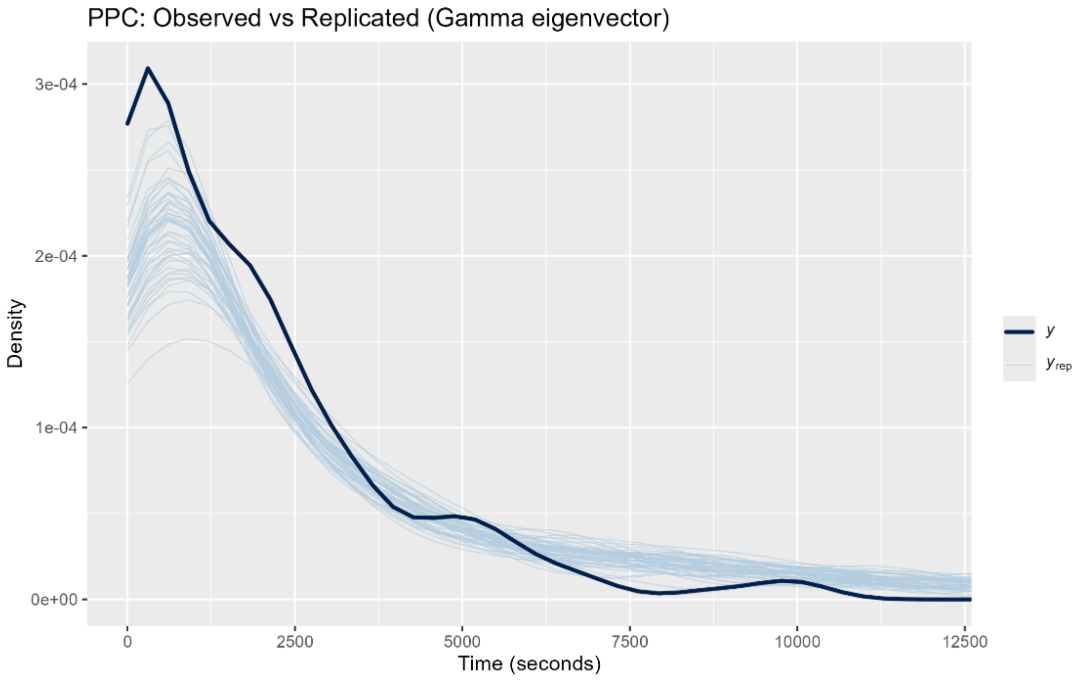
